# Computational Mesoscale Framework for Biological Clustering and Fractal Aggregation

**DOI:** 10.1101/2023.02.14.528441

**Authors:** Elnaz Zohravi, Nicolas Moreno, Marco Ellero

## Abstract

Complex hierarchical clustering mediated by diffusion and reaction is ubiquitous to many naturally occurring phenomena. The aggregates typically exhibit a fractal behavior or non-integer size scaling compared to their intrinsic dimensionality (2 – 3 dimensions). Such fractal aggregates have attracted attention in studying biological (i.e. bronchi and nervous system morphogenesis, blood clotting) and synthetic (i.e. colloids, polymers, catalysts, nano-dendrites, multicellular organisms) systems. In general, biological clustering can occur on a wide range of spatial/temporal scales, and depending on the type of interactions, multiple mechanisms (or stages) can be involved. As a consequence, the modeling of biological clustering is typically a challenging task, requiring the use of a variety of methods to capture the characteristic behavior of specific biological systems. Herein, we proposed a generalized-mesoscale-clustering (GMC) framework that incorporates hydrodynamic interactions, bonding, and surface tension effects. This framework allows for studying both static and dynamic states of cluster development. We showcase the framework using a variety of biological clustering mechanisms, and further illustrate its versatility to model different scales, focusing on blood-related clustering ranging from fibrin network formation to platelet aggregation. Besides the introduction of the mesoscale clustering framework, we show that a single biomarker (such as fractal dimension) is insufficient to fully characterize and distinguish different cluster structures (morphologies). To overcome this limitation, we propose a comprehensive characterization that relates the structural properties of the cluster using four key parameters, namely the fractal dimension, pore-scale diffusion, as well as the characteristic times for initiation and consolidation of the cluster. Additionally, we show that the GMC framework allows tracking of bond density providing another biomarker for cluster temporal evolution and final steady-state. Furthermore, this feature and built-in hydrodynamics interactions offer the potential to investigate cluster mechanical properties in a variety of biological systems.

## Introduction

Clustering, gelling, or coagulation are very common phenomena. Clusters are groupings of atoms, molecules, or ions that stick together due to a variety of physical-chemical interactions. They are important in physics [1, 2], chemistry [3, 4, 5], biology (i.e. multicellular organisms [6, 7], blood clotting [8], bronchi [9], nervous system morphogenesis [10] and tumor cells in cancer [11, 12]). Biological clustering is defined as the formation of higher-molecular-mass species as a result of the adhesion of smaller species. Different factors (e.g. species concentration, interactions, chain reactions, etc) can affect the clustering process leading to specific biological responses or functionality. For blood coagulation, for example, key factors that govern the clustering process have been identified: *i*) the concentration of biological species; *ii*) stages of coagulation; *iii*) characteristic time scales; *iv*) activation and aggregation of the species; and *v*) complex interactions including adhesion or linking. The existence of this multiplicity of governing factors poses significant challenges when modeling coagulation. Typical approaches involve the construction of system-specific models targeting narrower aspects of the coagulation cascade [8, 13]. However, comprehensive clustering models suitable for a wide range of factors and non-system specific are still missing.

A large part of the computational studies in clustering focus on investigating the fractal features of percolating clusters [14, 15, 16]. Examples of those schemes include diffusion-limited aggregation (DLA) [17, 18, 19, 20, 21, 15], ballistic aggregation (BA) [22] and reaction limited aggregation processes (RLA) [20]. It has been identified that cluster fractality (a.k.a fractal dimension *d_f_*) can provide relevant information about the physical, chemical, and biological properties of the systems [23, 24, 25]. Carlock et al [26], introduced a stochastic scheme that allows the formation of clusters with various morphologies and incorporates an effective interaction or aggregation range. Other numerical investigations using particle-based methods include Monte Carlo [27], molecular dynamics simulation [8, 28], and dissipative particle dynamics simulation [29, 30]. These methodologies have helped to reveal some of the fundamental aspects of clustering and showed the capability to reproduce diverse cluster fractality and morphology typically observed in aggregation processes. However, limitations on these approaches still exist since they target limited spatial-temporal scales (or even neglect temporal evolution), focusing mostly on steady-state features of the cluster. In general, due to the complexity of the dynamical process, a complete understanding of the factors that determine the evolution and steady state of the clusters remains hidden. This issue can be exacerbated when only one descriptor (i.e. *d_f_*) is used to rationalize the clustering. In the context of biological clustering, the number of studies that explore a range of particle interactions, initial conditions, system parameters, and spatial/temporal scales is still scarce.

Here, we present a generalized mesoscale clustering (GMC) framework that integrates hydrodynamic interactions, permanent bonding, and surface tension effects to model clustering in biological systems. To address the limitations of current methods in modeling relevant spatial-temporal scales, we employ the smoothed dissipative particle dynamics (SDPD) method [31, 32, 33, 34, 35]. The SDPD method discretizes the fluctuating Navier-Stokes equations and consistently satisfies the First and Second Laws of Thermodynamics and the Fluctuation-Dissipation Theorem [32, 34, 36, 37]. It has been used to study complex fluids, colloid-solvent interactions, colloid-colloid interactions, and blood flow [35, 38, 39, 40]. Our proposed model, based on the modified SDPD method [35, 31, 41], is tunable and versatile and reveals the correlation between the changes in cluster properties and physical model parameters. We demonstrate that initial conditions and interparticle interactions play a critical role in determining both static and dynamic behaviors of clusters.

We showcase the GMC framework using a variety of biological clustering mechanisms. To characterize the cluster’s evolution and steady state, we use *biomarkers* such as the number of bonds, fractal dimension (df), pore-scale diffusion (D), and time scales (*τ_I_* and *τ_G_*) of initiation and consolidation. Unlike previous models that prescribe the fractal dimension, our model allows for arbitrary values. Our results show that the use of multiple biomarkers is crucial for a correct characterization of the clusters, as aggregates with similar fractal dimensions may have different transport-related and mechanical properties. The paper details the cluster formation mechanisms, numerical schemes of the GMC framework, and the biomarkers definition, followed by results and discussion

## Clustering System Definition

We define three stages as fundamental ingredients for a biological clustering process, *activation, adhesion*, and *aggregation*. Additionally, we consider that the systems are constituted by three different types of species: *passive* (**P**), *active* (**A**), and *solvent* (**S**). Each species is represented in our methodology as an individual type of particle (see Fig. 1.*a*). Conceptually, this definition allows for the representation of clustering entities with different states of activity or reactivity. The change from passive to active states is referred to as *activation* stage. The existence of activity states and their transition is characteristic in the aggregation of particles in different systems such as platelet [42], fibrin (monomer/oligomer) [43], tissue factor [44], thrombin [45] in the blood coagulation system. For the initial condition of our system, we place randomly passive and solvent particles in the domain. The initial concentration of (**P**) particles is denoted as *ϕ_int_* = (*N_p_/N_t_*) where *N_p_* and *N_t_* are the numbers of passive and total particles in the system, respectively.

**Figure 1:**
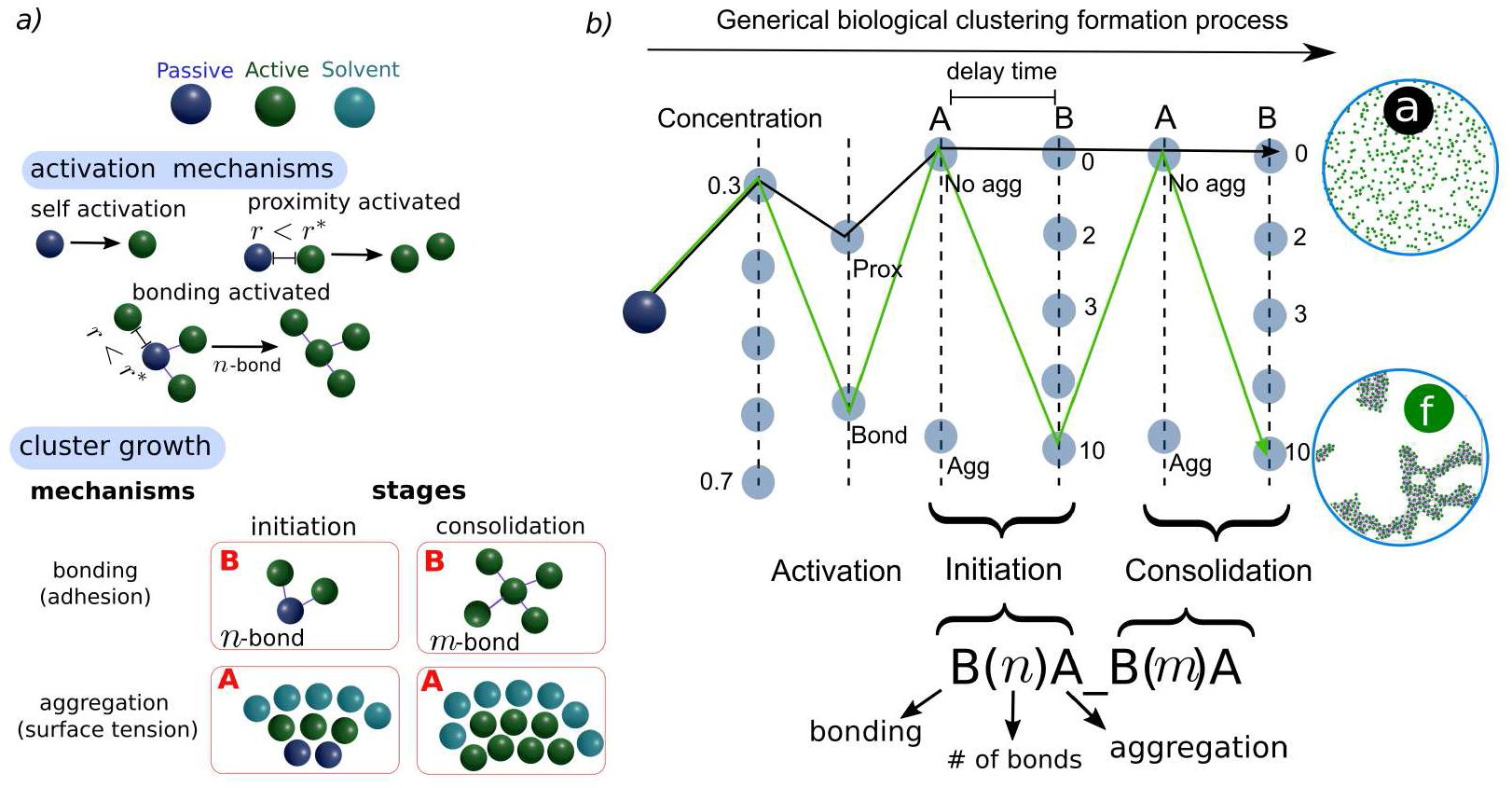
Schematic of the clustering system. *a*) Types of particles representing **(P), (A),** and **(S),** activation mechanisms proposed to capture a variety of biological activation processes, and cluster growth mechanisms. *b*) Generalized nodes diagram for clustering formation process and shorthand notation used to describe different clustering processes. Paths connecting nodes can be used to represent different clustering processes involving activation, initiation, and consolidation stages. Similarly, each stage can indicate the type of interaction (adhesion and/or aggregation). Examples of two paths (**a** and **f**) investigated herein are included for comparison.

### Activation

*Activation* of the species in the biological clusters can be mediated by complex chemical interactions with smaller molecules and other aggregating species. Activation proceeds at various rates depending on the mechanism, producing either free active or clustered active species [46]. The activation method, probability, and delay have a significant effect on the later growth of the cluster. Moreover, the activation can also determine the porosity, fractality, structure, and strength of the cluster [8, 6]. To model (**P**) particle activation, we proposed two activation schemes, denoted, proximity and bonding activation, as depicted in Fig. 1.*a*. The criteria used for each type of activation are as follows.

- Proximity activation: We define an activation-cutoff radius *r\**, such that, when the distance *r_pa_* between a (**P**) and an (**A**) particle is smaller than *r\**, the (**P**) particle becomes an (**A**) particle.
- Bonding activation: We define an activation-cutoff radius *r\** and a bond-number threshold *n*, such that, when the distance *r_pa_* between a (**P**) and an (**A**) particle is smaller than *r\**, a permanent bond is formed between them. The bonding formation process repeats with nearby (**A**) particles. When the number of bonds of the (**P**) particle reaches the threshold n for activation, the (**P**) particle becomes active.

In principle, the activation of a particle can be triggered as soon as the particle is located within the activation-cutoff radius. However, in many physical systems depending on biological conditions [43, 47], the activation may not take place immediately but with a certain probability or frequency. This translates in an effective *delay time* (*τ*_delay_) between the activation-triggering mechanism and the other existent particle interactions. Unless otherwise stated, in the simulation results presented, we initiate the clustering of the particles by placing a single (A) particle in the center of the domain, and *τ*_delay_ = 0.

Since biological clustering *via* permanent and transient interactions are quite prevalent in nature. The proposed framework is equipped with energy potentials (see Simulation methods section) to account for both types of driving forces. Depending, on the particular field, permanent interactions are associated with the strong-chemical bond formation and are usually referred to as *adhesion*. Transient interactions, in contrast, are generally weaker and long-ranged associations that allow for relative particle rearrangements. Multicellularity is a perfect example of these type of associations in biology [6]. Incomplete cell separation processes during cellular division lead to the formation of permanent bonds, ensuring that mother and daughter cells continue physically connected. This phenomenon occurs in both prokaryotes and eukaryotes (bacteria, algae, fungus, and at various phases of animal development) spanning numerous classes of multicellularity. [48, 49]. Reversible aggregations between cells have been also evidenced simultaneously or in combination with permanent bonds [50]. In transient interactions, the cells can either express sticky, velcro-like surface proteins that interact with proteins on the surfaces of neighboring cells, or they can excrete an extracellular viscoelastic matrix that holds them together. We describe the modeling of permanent and transient associations in the following sections.

### Aggregation growth

In the current study, aggregation is defined as the association of affine particles in a phase-separation process, as opposed to adhesion, which is defined as a permanent connection. Aggregation occurs because the unbalance in the affinity (interfacial tension) of the particles induces the formation of interfaces to minimize the energy of the system. To model the aggregation mechanism in our GMC, we use pair-wise forces to simulate the *affinity* between particles. Thus, depending on the type of particle, we can denote aggregation interactions as *A_αβ_*. where *α,β* = **P, A, S.** The extension of the proposed aggregation model can be readily used for studying biological clustering. Cellular and protein aggregation processes can lead to phase separation of the species, and the characterization of such systems in terms of their effective surface tension is already available in the literature [51].

### Adhesion growth stages

When fluid-like features (such as the aggregation previously discussed) are paired with elastic properties seen on shorter timescales or lower energy scales, viscoelastic type of structures are formed. Here, similar viscoelastic properties are modeled using permanent bonding forces between particles. To provide a generalized scheme able to reproduce a variety of biological clustering mechanisms we decompose the adhesion in two different consecutive stages, denoted as *initiation* and *consolidation* (see Fig. 1).*a*. A typical example of these stages is found in the blood coagulation cascade

- Initiation: The notion of initiation implies that particle bonding or proximity enables activation and a maximum number of n-bonds are formed between (**P**) and (**A**) before activation occurs. The definition of n-bonds can be correlated with biological properties by the specific type of interactions needed for a passive particle to be activated or even using a stoichiometric basis. This scheme imitates, for example, the initiation phase of the blood clotting cascade that is regulated by the activation of a number of clotting factors. It has been shown that binding of fibrin/fibrinogen to the integrin receptor results in activation cascades [52]. The characteristic time scale of this stage is denoted as *τ_I_*, and correspond to the time at which all the passive particle have been activated. For practical purposes, we consider that the initiation stage is concluded when at least 98% of the passive particles have changed to active.
- Consolidation: The consolidation (or maturation) stage of an individual particle is modeled considering that the maximum number of *m*-bonds can be created between (**A**) particles. The definition of *m*-bonds can be directly correlated with static and dynamic properties of the biological clusters, as the interconnectivity of the generated network and the strength of the associated bonds can be tuned. This stage is exemplified by the consolidation of platelet plugs during blood coagulation, where fibrin-mesh structures are formed [53]. The characteristic time scale of this stage is *τ_C_*. We must remark that for the whole system, the consolidation stage can overlap the initiation one since active particles can adhere to each other, while passive particles are still present. For physical systems, the degree of overlapping between stages can be a characteristic indication of the reactivity of (**P**) particles, and the mechanical evolution of the cluster.

Depending on the physical system, it is customary to associate the gel point to the condition where the system undergoes a change in viscosity due to the formation of an interconnected network. In general, since the connectivity of the network can keep evolving during the consolidation stage, here we define the gelling time *τ_G_* as the condition where the rate of the bond formation reaches a steady condition. For practical purposes, in our simulations, we consider that steady condition when the change in the number of bonds between 10^4^ time steps is lower than 0.1%. Therefore, the consolidation stage can in principle continue after gelling occurs. Similar descriptions of *τ_G_* can be found in biological clustering where the complete network is not established at the gel point, and new fibers and branch points can form afterward [54].

### Cluster formation mechanisms

To facilitate the discussion of the results for the different clustering schemes, we introduce the following notation. Initiation and consolidation stages of adhesion are described in terms of the driving force used, bonding *B*(*k*), and aggregation is described as *A*, where *k* denotes the number of bonds threshold for the stage. Thus, a stage where bonding and aggregation occur at the same stage is expressed as *B*(*n*)*A*, whereas a stage with only bonding can be simply represented with *B*(*n*). The whole clustering mechanism can be written in compact notation as *B*(*n*)*A_B*(*m*)*A*. Here, *n* and *m* correspond to the threshold number of bonds in the initiation and consolidation stages, respectively. Given the generality of the framework and the possibility to explore different parameters and cluster mechanisms (type of activation, stages with a varied number of bonds, or use of aggregation), the number of possible combinations to showcase the GMC is significantly large. Thus, in Fig. 1.*b*, we introduce a generalized nodes diagram for the clustering formation process. In this diagram, a series of vertical parallel lines are used to indicate relevant factors such as (**P**) particle concentration, activation mechanism, bonding number, and aggregation for the different stages. Discrete nodes along each factor indicate the specific value adopted, and the path connecting those nodes fully describes the clustering process.

For simplicity, we will streamline the presentation of the GMC framework and the cluster characterization using 7 representative mechanisms. These mechanisms are chosen to illustrate different aspects of the methodology proposed and highlight the need for complementary biomarkers for cluster analysis. These mechanisms are denoted as:

- *a*) Proximity Activated (*Pr*)
- *b*) Proximity Activated-Aggregation (*Pr_A*)
- *c*) Bonding Activation (*B*(3)_*B*(0))
- *d*) Bonding Activation (*B*(3)_*B*(2))
- *e*) Bonding Activation (*B*(3)_*B*(10))
- *f*) Bonding Activation (*B*(10)_*B*(10))
- *g*) Aggregation-Delay Bonding Activation (*B*(3)*A_ps_B_*(0))

Fig. 1.*b*, depicts the whole path for mechanisms of *(a)* and *(f)* at *ϕ_int_* = 30%. In SI Fig. S2 we present the corresponding paths for all the mechanisms *(a)*–*(g)*. Fractal dimension and pore-scale transport analysis are performed for mechanisms *(a)-(f)*, whereas mechanism *(g)* is defined to explore activation-delay time (*τ*_delay_) on the evolution of clusters where phase separation (aggregation) is a dominating effect over adhesion (bonding). To this end, different *τ*_delay_ between bonding events and aggregation are explored.

In the results section, we investigate how the morphology and other properties of the clusters are determined by different factors that define the system. The factors of interest are: *i*) the initial concentration (*ϕ_int_*) of (**P**), ii) the maximum number of bonds *n* and *m* (during initiation and consolidation), and *iii*) the balance between aggregation and adhesion (bonding) phenomena. To explore (*i*), we focus on medium to larger concentrations relevant to many biological applications. In particular, we simulate the range *ϕ_int_* = [20%, 30%, 40%, 50%, 60%, 70%]. In biological cases, such as blood coagulation, the concentration of the aggregating species is a key factor that determines the ultimate qualities of clots [55, 56]. The broad range of concentrations used herein may have potential applications to investigate clustering in a variety of circumstances, such as clot formation in healthy and unhealthy patients. Regarding (*ii*), the maximum number of *n* and *m* bonds are crucial for the connectivity and form of the clusters. In physical applications, a reduction in fiber diameters is typically connected with an increase in the number of branch points in the aggregate [57]. Factor (*iii*), is relevant in the clustering process where the strength of the aggregation and adhesion can vary dynamically depending on physiological conditions. For example, at high-shear stresses platelets can dominantly associate *via* aggregation, whereas firm attachment mediated specific receptors may occur in a later stage, inducing an activation delay[43, 47, 58].

## Simulation methods

We now describe the mathematical models that describe the hydrodynamic interactions, aggregation, and adhesion, between the different (**P**), (**A**), and (**S**) particles.

### Mathematical model for hydrodynamic interactions

We describe the mass and momentum balance of our system in terms of the Navier-Stokes equations. In a Lagrangian reference frame, the mass balance is given by *dρ/dt* = ∇ · **v**, and the momentum equation reads *ρ*d**v**/*dt* + ∇*p* – *η*∇^2^**v** – (*ζ + η/D*) ∇∇ · **v** = **f**^external^ for *D* = 2, 3 (1), where *p* is the pressure, whereas *η* and *ζ* are the standard shear and bulk viscosity.

We define the mass and momentum balance using the SDPD method such that consistent thermal fluctuations are incorporated. Using this description, we can model the transport of the different particles in the systems at mesoscales. The evolution equations for the position of the particle is *d***r***_i_/dt* = **v***_i_*, whereas the discrete density d*_i_* and the deterministic part of the momentum Eq. (1), without external forces, can be expressed as

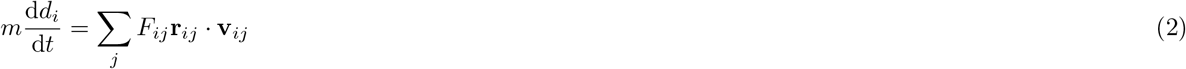

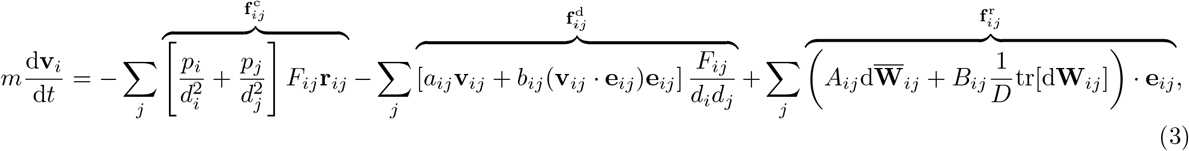

where **v***_ij_* = **v***_i_* – **v***_j_, a_ij_* and *b_ij_* are friction coefficients related to the shear *η* and bulk *ζ* viscosities of the fluid through *a_ij_* = (*D* + 2)*η/D* – *ζ* and *b_ij_* = (*D* + 2)(*ζ* + *η/D*). *D* is the dimension of the system. The last term in (3), consistently incorporates thermal fluctuations in the momentum balance. A complete description of the terms *A_ij_, B_ij_*, the equation of state used to define the pressure *p*, and the form of the function *F_ij_* are given in SI Eqs.(S1-S5).

### Mathematical models of aggregation

We incorporate the interfacial tension effects between the different types of particles (**P**, **A**, and **S**) by including in the momentum equation (3) an additional pair-wise force *F*^int^ [59, 60, 41], such that 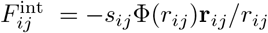 (4) where

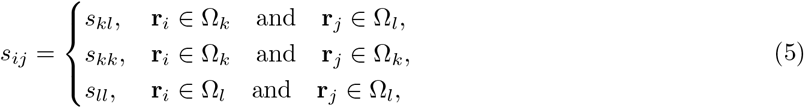

and Φ(*r_ij_*) is a shape factor given by 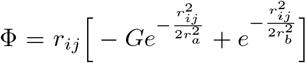, where *G* = 2^*D*+1^, being *D* the dimension. The range for repulsive and attractive interactions is defined as 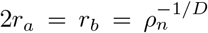, such that a relatively uniform particle distribution is obtained for a given surface tension *σ*. The interaction parameters satisfy *s_kk_* = *s_ll_* = 10^3^*s_kl_*, and the magnitude can be obtained from the surface tension and particle density of the system as [60]

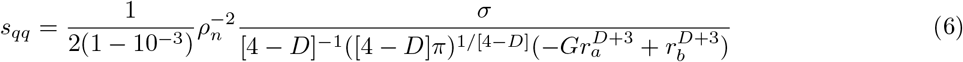

### Mathematical models of adhesion

The mathematical model of adhesion in this work is modeled by Morse bonding potential [61] as this equation, 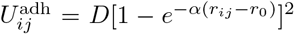 (7). Among the several molecular potentials, the Morse potential is an ideal and typical anharmonic potential that explicitly takes bond-breaking effects into account, which is useful for our future investigations, leading to an adhesion force 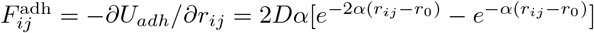 (8).

### Simulation details

Without loss of generality, we focus our investigations on two-dimensional systems and consequently reported clusters with characteristic *d_f_* ranging from 1.4±0.01 to 2±0.01. We use a 2D periodic box for simulations. The characteristic parameters of the SDPD method are shown in Table. S1 and S2 display values relating to surface tension and permanent bonding potentials. To account for the influence of thermal noise and the intrinsic randomness of the clustering process, all the simulations were conducted over ten randomly generated particle positions and velocities. All the simulations were conducted using a modified version of the open-source software LAMMPS [62, 63].

### Biomarkers definition

We define a set of four biomarkers to characterize both the static and dynamic process of cluster formation. The biomarkers are the fractal dimension (*d_f_*), initiation time (*τ_I_*), gelling time (*τ_G_*), and the pore-scale diffusion 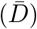, given by the mean-squared-displacement (MSD) of (**S**) particles after cluster consolidation. The proposed biomarkers have been independently used as relevant experimental parameters for cluster characterization [64]. For example, fibrous proteins such as fibrin can form a complex network or gel structures typically characterized by *d_f_* and gel point analysis [65]. The gelling time is widely used in clinical tests to show improper biological functionality. Even though, these two biomarkers are tightly linked [66] the entire network is rarely created at the gel point, with extra fibers and branch points added later [54]. As a consequence, there has been evidence of the relevance to adopt more than one biomarker for proper cluster characterization. Regarding the pore-scale diffusion effects, they provide a sensitive measure of pore size and total cluster shape. In general, the structure of percolated clusters has little effect on the diffusion of molecules much smaller than their pore size. However, the perfusion of fluids [67] or nanoparticles through their structure can exhibit characteristic hydrodynamic interactions that lead to anomalous diffusion.

Other properties used for cluster characterization include fiber diameter, density, number and nature of branch points, and distances between branch points. Similarly, here we track during the simulations the evolution in the number of bonds between active particles (*N_B_*), active particles number (*N_A_*), passive particles number (*N_P_*), and cluster’s radius of gyration (*R_g_*). We calculate the radius of gyration *R_g_* of a cluster using 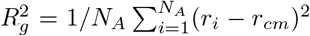 (9), where *r_i_* is the position of ith (**A**) particle and *r_cm_* is the center of mass of the cluster.

#### Fractal dimension evaluation

It is common in the analysis of clusters to calculate *d_f_* from the radius of gyration, *R_g_*, using a linear fit to *R_g_* = *kN^β^* on a log-log scale, where *k* is a constant, N is the number of aggregated particles, and *d_f_* = 1/*β* is the estimated fractal dimension. As long as the number of particles in the cluster is high, it is expected that the slope remains approximately constant. The radius of gyration algorithm includes randomly selecting a sufficient number of particles and computing *R_g_* of the selected particles according to their center of mass. We measure *d_f_* in both the cluster’s steady state and during its time evolution.

#### Fluid mean-squared displacement evaluation

Cellular transport is reduced in biological environments due to the presence of macromolecules, such as the trapping of cells in clots of fibrin [64, 68], or nutrients and toxins diffusion in mycelial networks [69]. Anomalous diffusion is a key theoretical scenario that explains molecular transport in complex settings and the cause for sub-diffusion, in which the solute is not allowed to occupy a certain amount of space. Here we use the mean-squared displacement (MSD) to investigate the diffusion of (S) particles when the clusters have attained a steady state or gel point. See SI. 2 for a detailed description of the MSD calculation.

SI Fig. S1 shows a schematic representation of MSD as a function of time for pure normal diffusion and transient anomalous subdiffusion (AD) in the complex clusters. AD exists between 2 crossover times (*t*_*CR*1_ and *t*_*CR*2_). When *t* < *t*_*CR*1_, or at very short diffusion times, normal diffusion is present and is defined by *D* = *D*_0_. however, when *t* > *t*_*CR*1_, or at long diffusion times, normal diffusion is still present and is defined by a finite value of *D* = *D*_∞_. The anomalous diffusion exponent measures the deviation from normal diffusion. Diffusion is Gaussian at long diffusion times according to the central limit theorem, with a finite effective diffusivity 0 < *D*_∞_ < *D*_0_ [70].

## Results

We consider specific microscopic parameters (concentration and mechanism type) and study how they affect the above-mentioned biomarkers. We perform our simulations for concentrations ranging from *ϕ_int_* = 20%, 30%, 40%, 50%, 60%, 70%, and the six mechanisms described previously (see also SI Table. S3). To streamline the discussion, we select a reduced data set of results that highlight the main findings. An extended breakdown of the structures obtained and their biomarkers for the whole range of systems is included in the Supplementary Information.

### Effect of concentration and mechanism type on *d_f_*

In this section, we compute the value of *d_f_* for various initial concentrations of (P) particles, *ϕ_int_*, and mechanism types. SI Table. S3, summarizes the *d_f_* values, with a standard deviation of ±0.01, for initial concentrations of *ϕ_int_* = 30%, 40%, 50% and the six mechanisms evaluated. Fig. 2 illustrates the morphology of *ϕ_int_* = 40% for different mechanisms, while SI Figs. (S3, S4) for higher and lower concentrations.

**Figure 2:**
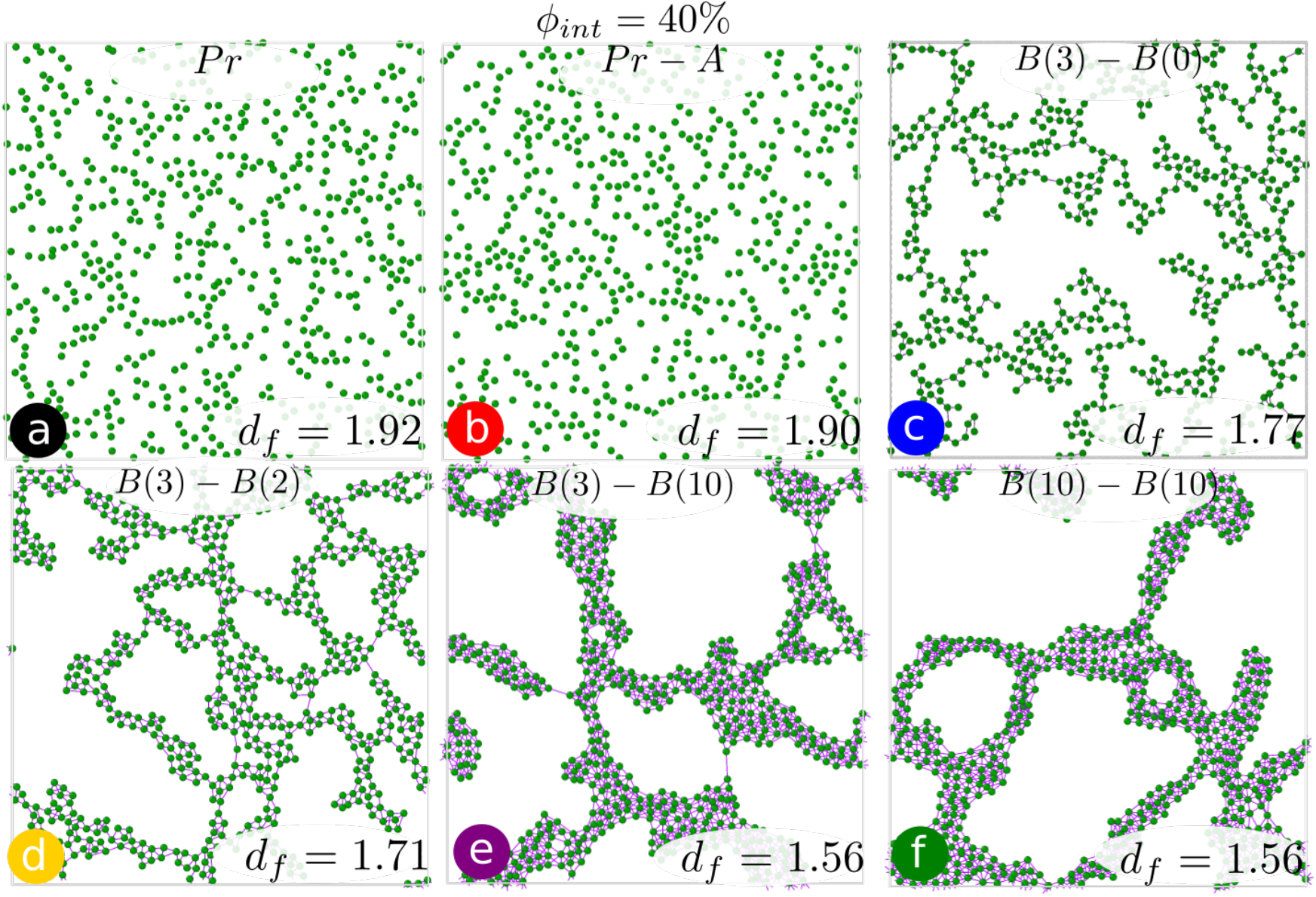
Final cluster morphology and *d_f_* are shown for six alternative mechanisms (*a, b, c, d, e, f*) and *ϕ_int_* = 40%

In general, we observe that the interplay between passive particle transport, bonding, and aggregation plays a crucial role in determining the magnitude and sensitivity of *d_f_*. We found that *d_f_* increases with the concentration of (P) particles, *ϕ_int_*. For concentrations greater than 50%, *d_f_* ~ 2 (indicating space-filling in 2D), and the magnitude of *d_f_* becomes insensitive to variations in clustering mechanisms. This is because the probability for particle encounters scales with the concentration, leading to the rapid formation of clusters that occupy the entire space and reducing the influence of particle-transport effects. The increase in *d_f_* with concentration is attributed to the ability of more particles to diffuse into the internal regions of the forming aggregate before being irreversibly bound and the ability of aggregates to interpenetrate before bonding occurs.

In contrast, at lower concentrations (Fig. 2), both particle-transport and particle interaction effects are relevant in shaping the diversity of *d_f_*. Lower values of *d_f_* are observed as the number of bonds increases. Additionally, this reduction is significantly sensitive to bonding during the consolidation stage. Bonding-mediated activation results in the preliminary formation of highly-branched clusters with closed pores. However, as activation proceeds, particle mobility is reduced, limiting their ability to penetrate the internal regions of a growing cluster. As a result, the sensitivity of *d_f_* with the number of bonds (*n*) in the activation stage exhibits a threshold, beyond which further increase in *n* does not affect the fractality of the sample (see Fig. 2 for mechanisms *e* and *f*). Further reduction in *d_f_* can be achieved in clustering processes that exhibit a consolidation stage. The reduced mobility due to activation is overcome by local rearrangements of the particles, leading to dense clusters with large closed pores. However, we must note that the sensitivity of *d_f_* with the number of bonds (*m*) during the consolidation stage is limited by steric hindrance.

### Effect of concentration and mechanism type on MSD of (S) particles

This section computes MSD values for various initial concentrations of (P) particles, *ϕ_int_*, and mechanism type. To compute the MSD of **(S)** particles, we first freeze the cluster particles or **(A)** particles while freely moving (S) particles for 10^6^ time steps (a long enough period of time). (S) particles do not penetrate the clusters but only diffuse in the existent pores. MSDs are computed as an ensemble average over solvent particles.

#### Effect of concentration on anomalous diffusion exponent, *α* and *D*_∞_

Fig. 3.*a* shows the mean time-averaged MSD as a function of lag time (log-log scale) for *ϕ_int_* = 20%, 30%, 40%, 50%, 60% and mechanism *c* (*B*(3)_*B*(0)). For comparison, we include the corresponding MSD for pure solvent particles (unconfined). In Fig. 3, normal diffusion for pure fluid (*ϕ* = 0%) exhibit a characteristic slope, *α* = 1, whereas for anomalous diffusion *α* ≤ 1. In general, we observe that anomalous diffusion emerges at short-to-intermecliate time scales, while at longer time scales normal diffusion is restored, reaching the limiting value *D*_∞_. The transition from anomalous to normal diffusion for each system occurs at the characteristic crossover time *t*_*CR*2_.

**Figure 3:**
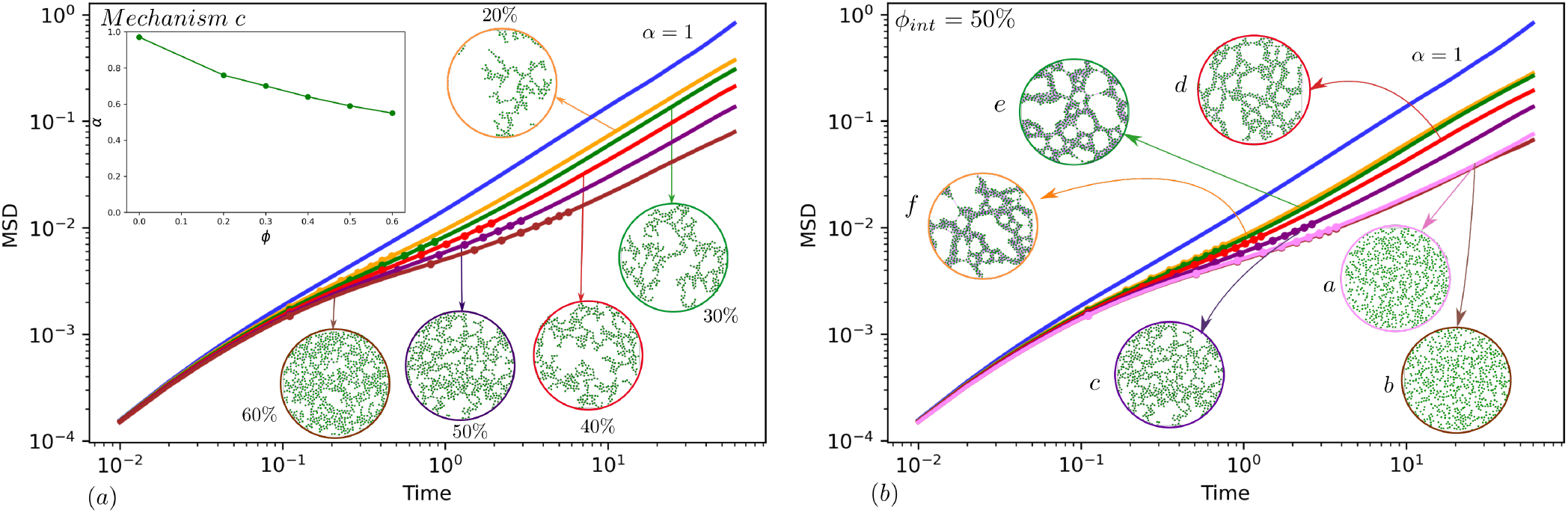
a) Mean time-averaged MSD as a function of lag time (log-log scale) for mechanism *c, ϕ_int_* = 20%, 30%, 40%, 50%, 60%, along with pure fluid. Solid markers indicate the range of anomalous diffusion, b) Mean time-averaged MSD as a function of lag time (log-log scale) for *ϕ_int_* = 50%, mechanisms *a, b, c, d, e* and *f* along with pure fluid. Final cluster morphology is shown for each curve.

As expected, we identify that *α, t*_*CR*2_, and *D*_∞_ are all functions of cluster concentration. In Table. 1, we give a breakdown of the non-dimensional diffusion coefficient 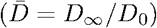 and *α* for different concentrations and mechanisms. Where *D*_0_ is the diffusion coefficient of the pure solvent. The deviation of MSD from normal diffusion grows as cluster concentration increases, indicating a consistent deviation from Fick’s normal diffusion towards a sub-diffusion. The insight plot in Fig. 3.*a* depicts a smooth reduction of *α* with the concentration, due to the increment in the confinement of the solvent particles. Overall, we find that as the concentration increases, the crossover time increases (see solid markers in Fig. 3), whereas *α* and *D*_∞_ decreases.

**Table 1:** Non-dimensional Diffusion Coefficient 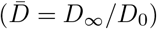 and anomalous *α* for different initial concentration of (**P**) particles, *ϕ_int_* and different mechanisms. The standard deviation of *α* and 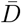 is equal to ±0.001

| Mechanisms | Concentrations |  |  |
| --- | --- | --- | --- |
| | $\phi_{int} = 30\%$ | $\phi_{int} = 40\%$ | $\phi_{int} = 50\%$ |
| *a* | $\alpha = 0.64, \bar{D} = 0.36, d_f = 1.84$ | $\alpha = 0.58, \bar{D} = 0.29, d_f = 1.92$ | $\alpha = 0.52, \bar{D} = 0.25, d_f = 1.98$ |
| *b* | $\alpha = 0.65, \bar{D} = 0.44, d_f = 1.81$ | $\alpha = 0.59, \bar{D} = 0.31, d_f = 1.90$ | $\alpha = 0.53, \bar{D} = 0.27, d_f = 1.96$ |
| *c* | $\alpha = 0.70, \bar{D} = 0.55, d_f = 1.65$ | $\alpha = 0.64, \bar{D} = 0.42, d_f = 1.77$ | $\alpha = 0.59, \bar{D} = 0.32, d_f = 1.87$ |
| *d* | $\alpha = 0.74, \bar{D} = 0.62, d_f = 1.48$ | $\alpha = 0.68, \bar{D} = 0.55, d_f = 1.71$ | $\alpha = 0.65, \bar{D} = 0.50, d_f = 1.82$ |
| *e* | $\alpha = 0.79, \bar{D} = 0.78, d_f = 1.43$ | $\alpha = 0.75, \bar{D} = 0.68, d_f = 1.56$ | $\alpha = 0.69, \bar{D} = 0.62, d_f = 1.75$ |
| *f* | $\alpha = 0.79, \bar{D} = 0.79, d_f = 1.43$ | $\alpha = 0.75, \bar{D} = 0.69, d_f = 1.56$ | $\alpha = 0.71, \bar{D} = 0.63, d_f = 1.75$ |

The subdiffusive behavior observed is due to the dynamics of particles trapped inside the pores. At high concentrations, a significant portion of particles is confined in closed pores, resulting in restrictions on their motion and oscillations of individual particle trajectories that follow Gaussian statistics. However, since not all particles have the same degree of confinement, the power-law (*α* = 1) dependency can be regained for long time scales, when the MSD is averaged over all particles, including less confined ones. Nevertheless, individual particles may still exhibit subdiffusive behavior. Understanding the distribution of diffusive behaviors among particles and its effect on the ensemble average of MSD is a vital aspect of characterizing diffusion in dense clusters and confined systems. [70]

#### Effect of mechanism type on anomalous diffusion exponent, *α* and *D*_∞_

Fig. 3.*b* shows mean time-averaged MSD as a function of lag time (log-log scale) for *ϕ_int_* = 50% and six mechanisms to evidence how aggregation, bonding in both stages of initiation and consolidation affect the magnitude and sensitivity of MSD values. In general, we find that *α, D*_∞_, and *t*_*CR*2_ are highly sensitive to the activation type. Values of *α* and D∞ for proximity (a) and aggregation (b) activation mechanisms, are the lowest among all mechanisms, whereas the *t*_*CR*2_ value is the highest. These mechanisms are responsible for percolated clusters with reduced pore size. In contrast, when activation occurs via bonding mechanism (c,d), branched clusters with larger pore sizes lead to larger values of *α* and *D*∞. Further increase of *α* and *D*_∞_ is identified for the clustering mechanism that involves a consolidation stage (*e, f*), inducing the local rearrangement of the cluster particles leading to branched and more compact structures with increased pore size.

Our results demonstrate the impact of aggregation and bonding mechanisms on particle transport in (**S**). Different cluster structures result in varying subdiffusive motions. To further understand these findings, we compare *d_f_* values for different cases in Table. 1 and observe how some cases have similar *d_f_* but different 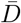 and *α* values. For example, the case with *ϕ_int_* = 50% and mechanism *f* has a *d_f_* value of 1.75, which is similar to the case with *ϕ_int_* = 40% and mechanism c (with *d_f_* = 1.77), but the values of 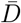 and *α* are different. This highlights the limitations of using *d_f_* as the sole criterion for clustering analysis. We suggest that a combined analysis of *d_f_* and MSD of the percolating fluid (or probes) provides a comprehensive characterization of the cluster. Furthermore, different MSD-related parameters such as *α, D*_∞_, and *t*_*CR*2_ offer important insights into biological systems, which are not revealed by *d_f_* alone. These parameters have been extensively studied in relation to biological systems [71, 70, 72, 73, 69]. Additionally, it is crucial to understand molecular transport in complex biological systems, where physical factors can impact biological processes. Different objects such as proteins bound or fibrin networks can have varying dynamics, which affect kinetic rates and structure. Our findings can be a starting point for further research toward more accurate models of congested biological systems.

### Effect of concentration and mechanism types on initiation time, *τ_I_* and gelling time, *τ_G_*

In Table 2, we summarize the measured gelling time (*τ_G_*) and initiation time (*τ_I_*) for various concentrations and mechanisms. To make the time scales dimensionless, we normalize *τ_I_* and *τ_G_* using the diffusive time (*τ_diff_*) as 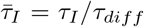 and 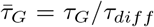. The diffusive time is calculated as *τ_diff_* = (4*h*)^2^/*D*_0_, where *h* is the kernel cutoff radius, which approximates the size of the SDPD fluid particle [74], and *D*_0_ is the diffusion coefficient of the pure fluid.

**Table 2:** *τ_G_* (gelling time) and, *τ_I_* (initiation time) for different initial concentration of (**P**) particles,*ϕ_int_* and mechanisms

| Mechanisms | Concentrations |  |  |
| --- | --- | --- | --- |
| | $\phi_{int} = 30\%$ | $\phi_{int} = 40\%$ | $\phi_{int} = 50\%$ |
| *a* | $\bar{\tau}_I = \bar{\tau}_G = 0.38$ | $\bar{\tau}_I = \bar{\tau}_G = 0.19$ | $\bar{\tau}_I = \bar{\tau}_G = 0.05$ |
| *b* | $\bar{\tau}_I = \bar{\tau}_G = 0.33$ | $\bar{\tau}_I = \bar{\tau}_G = 0.06$ | $\bar{\tau}_I = \bar{\tau}_G = 0.01$ |
| *c* | $\bar{\tau}_I = 1.48, \bar{\tau}_G = 1.63$ | $\bar{\tau}_I = 0.21, \bar{\tau}_G = 0.22$ | $\bar{\tau}_I = 0.04, \bar{\tau}_G = 0.05$ |
| *d* | $\bar{\tau}_I = 1.51, \bar{\tau}_G = 2.55$ | $\bar{\tau}_I = 0.26, \bar{\tau}_G = 1.12$ | $\bar{\tau}_I = 0.04, \bar{\tau}_G = 0.84$ |
| *e* | $\bar{\tau}_I = 1.55, \bar{\tau}_G = 6.46$ | $\bar{\tau}_I = 0.11, \bar{\tau}_G = 5.34$ | $\bar{\tau}_I = 0.03, \bar{\tau}_G = 5.11$ |
| *f* | $\bar{\tau}_I = 5.1, \bar{\tau}_G = 6.7$ | $\bar{\tau}_I = 1.63, \bar{\tau}_G = 5.35$ | $\bar{\tau}_I = 0.11, \bar{\tau}_G = 5.11$ |

Our results show that *τ_I_* and *τ_G_* have an inverse relationship with concentration, and this trend holds true for all mechanism types. Higher concentrations lead to faster consolidation or gelling due to the increased probability of particle interactions. For instance, an increase in platelet concentration at the site of injury increases the chance of platelet-platelet interactions, resulting in faster clot formation.

Regarding the type of mechanism, we find that the initiation time, *τ_I_*, and gelling time, *τ_G_* are the lowest for clusters due to the proximity (a) and aggregation (b) mechanisms, and the rise in those that activation is driven mainly by the bonding (c,d,e,f). Overall, we observe that the initiation time scale, *τ_I_*, is related to the number of first-stage bonds, whereas the gelling time scale, *τ_G_*, is linked to the number of second-stage bonds and increases with their number. Thus, by adjusting the number of bonds, we can regulate and tune these time scales.

Different gelling mechanisms can have varying kinetics, and this can impact the gelling time. For example, mechanisms that involve the formation of physical bonds, such as hydrogen bonding or van der Waals forces, tend to have slower kinetics and result in longer gelling times. On the other hand, mechanisms that involve chemical reactions, like the cross-linking of polymers, can result in faster gelation due to the rapid formation of chemical bonds. Understanding the relationship between initiation time and activation mechanism is important for comprehending the regulation of biological processes.

### Cluster evolution over time

#### Time evolution of several biomarkers

In this section, we focus our attention on the temporal evolution of the different biomarkers, to illustrate additional dynamical features that can be accounted for in the GMC framework. In Fig. 4 we compare the evolution *d_f_*, active particles number, passive particles number, the radius of gyration of the cluster, and bonds number for three mechanism types *c*) (*B*(3)_*B*(0)); *f*) (*B*(10)_*B*(10)); *g*) (*B*(3)*A_ps__B*(0)), at *ϕ_int_* = 40%. This allows us to explore the effect of surface tension (aggregation) between mechanisms *c* and *g*, and the consolidation effect between mechanisms c and f.

**Figure 4:**
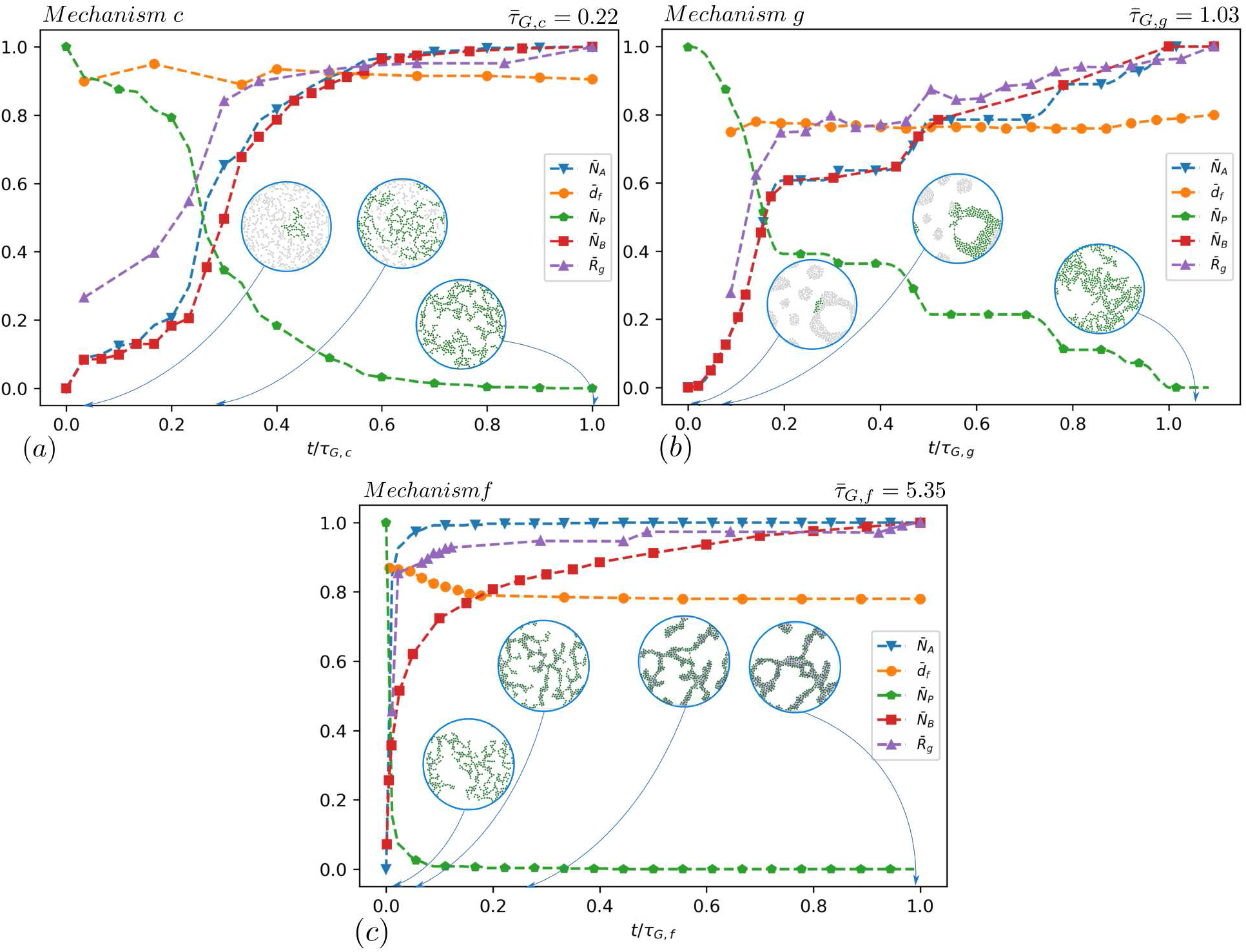
Time evolution of bond numbers, *d_f_*, active particle numbers, passive particle numbers and the radius of gyration of the cluster for *ϕ_int_* = 40%, a) Mechanism (*c*), b) Mechanism (*g*), c) Mechanism (*f*). Time value is made dimensionless with the *τ_G_* of each mechanism. Some intermediate cluster structures are shown during the gelling process for each mechanism.

In Fig. 4, we present the dimensionless values for the number of (**P**) particles, bonds, a radius of gyration, (**A**) particles, and fractal dimension. The non-dimensionalization is done using the corresponding steady-state values for each parameter, whereas for *d_f_*, we use the dimensionality of the box (2). The dimensionless values are denoted using the upper bar notation. The time scale is normalized using the gelling time, *τ_G_*, of each mechanism.

It can be observed that the fractal dimension *d_f_* changes as the cluster grows in all three mechanisms. Unlike traditional methods that keep the fractality constant during growth, our method shows a change in *d_f_* values over time. Taking as an example blood clotting phenomena, the evolution of the fractal dimension of a blood clot refers to the changes in the complexity of the shape of the clot and branching of the fibrin fibers over time and can change due to various factors such as the accumulation of additional platelets and other blood components. As the clot matures, the fractal dimension may stabilize.

In addition, we see a decreasing trend in (**P**) particles and an increasing trend in (**A**) particles, bond numbers, and radius of gyration. This can be correlated again with typical blood clotting stages, where an increase in active blood components, such as platelets, can lead to a more stable and stronger clot, while a decrease in passive blood components, such as plasma proteins, can weaken the clot and increase the risk of lysis. Understanding these changes in the fractal dimension, active and passive particle numbers, and bond numbers can provide important insights into biological cluster formation and behavior, and has potential applications in diagnosing and treatment of related diseases.

Mechanism *g* is introduced to investigate the effect of activation delay time on cluster morphology. During this process, there is a delay between activation (bonding) and aggregation events. The activation delay time reflects how the cluster’s evolution changes as aggregation takes precedence over bonding. In Fig. 4.*b*, we can observe a step-wise evolution of the biomarkers in mechanism *g*, indicating a sudden activation of passive particles that were previously together as a droplet. Compared to mechanism *c*, the fractal dimension of mechanism *g* is consistently smaller during the whole process, and the gelling time is approximately five times higher. The activation delay time is an important feature that characterizes also biological clustering in blood. It refers to the time interval between the start of a clotting event and the beginning of the active phase, during which platelets and clotting factors become activated. This time interval can impact the structure and stability of a blood clot and may affect clot formation and resolution. A longer activation delay time may result in a less stable clot with a simpler structure, while a shorter activation delay time may result in a more stable clot with a more complex structure.

The impact of the consolidation stage (second-stage of bonding) on the time evolution of biomarkers can be seen by comparing mechanisms *c* and *f* in Fig. 4.c. Mechanism *f* has a continuous formation of bonds, causing the structure of the cluster to continue evolving, even after converting (**P**) particles into (**A**) particles. As a result, the *d_f_* value of the cluster decreases over time, even after gelling, and is lower than mechanism c. The time gap between initiation and gelling is also larger in mechanism *f* compared to mechanism c, with a gelling time that is roughly 25 times higher. The consolidation stage as described by mechanism *f* resembles the final stages of blood coagulation, where the clot gradually solidifies, connects, and becomes more stable over time, changing from a loose and weak structure to a more organized and stable one. However, the exact timeline of this process can vary greatly, taking several hours to several days, depending on various factors.

#### Time evolution of bond numbers in various mechanisms

This section compares the time evolution of bond numbers in various mechanisms. The time evolution of bond number, 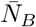, for mechanisms *c, d, e, f* and *ϕ_int_* = 40% is shown in SI Fig. S10. Bond numbers are dimensionless with their values in a steady state or gelling state, time values are also made dimensionless using the *τ_G_* of mechanism *f*. The changing bond number over time demonstrates that our method can alter the system’s kinetic properties over time in a manner similar to real biological systems. The time required to reach the steady state increases as the number of bonds in the second-stage bonding, *m*, increases. This allows us to construct clusters with a wide range of gelling times (depending on activation and aggregation mechanism conditions) and different final cluster structures.

Overall, we have shown that the interplay between activation, consolidation, and surface tension effects originate a multiplicity of transitions that mimic naturally occurring clustering processes. However, cluster morphology can be also marginally altered by systematic variation of the parameters in our model. This is the number of bonds allowed per stage, or variation on the delay time. Such systematic variation will be typically associated with the specific biological system modeled, and require an appropriate fine-tuning of the model parameters that are out of the scope of this study. However, in SI. 3.1, 3.2 and 3.3 we illustrate a preliminary case of study, where parametric variations in the bonding for different stages leads to different quantitative and qualitative results on cluster morphology.

## Discussions and Conclusion

In this study, we introduce a new method for modeling cluster growth based on the SDPD method, that incorporates hydrodynamics, adhesion-, and aggregation-type interactions between constituents. Our method can be used to study the effects of different characteristic parameters, such as initial concentration, and activation methods (proximity and bonding). Additionally, we could examine the effects of the first-stage bonding, second-stage bonding, and the delay time between the first-stage bonding (activation) and aggregation. The paradigm systematically identifies distinctive biomarkers such as fractal dimension (df), pore-scale transport properties (anomalous diffusion exponent, *α*, and infinite diffusion coefficient, *D*_∞_), and time scales (initiation time, *τ_I_*, and gelling time, *τ_G_*).

We summarise the results obtained with the GMC scheme in Fig. 5 for *ϕ_int_* = 40% (see SI Figs. S11, S12 for *ϕ_int_* = 30% and *ϕ_int_* = 50%). In Fig. 5, we present a *biomarker diagram* that displays the variety of clusters that can be generated and their corresponding characteristics. The biomarker diagram helps us to navigate the effects of concentration and clustering paths into the identified biomarkers, along with the cluster morphology. In Fig. 5, we show activation of a passive particle occurs in two ways: 1-Proximity and 2-Bonding. We define two mechanism types *a* and *b* for the proximity activation approach, in which mechanism *b* is combined with aggregation. The bonding approach has five mechanism types *c* – *g* with varying bonds number during activation and consolidation. In mechanism *g*, there is also aggregation, which has a *τ*_delay_ between aggregation and activation (first-stage bonding).

**Figure 5:**
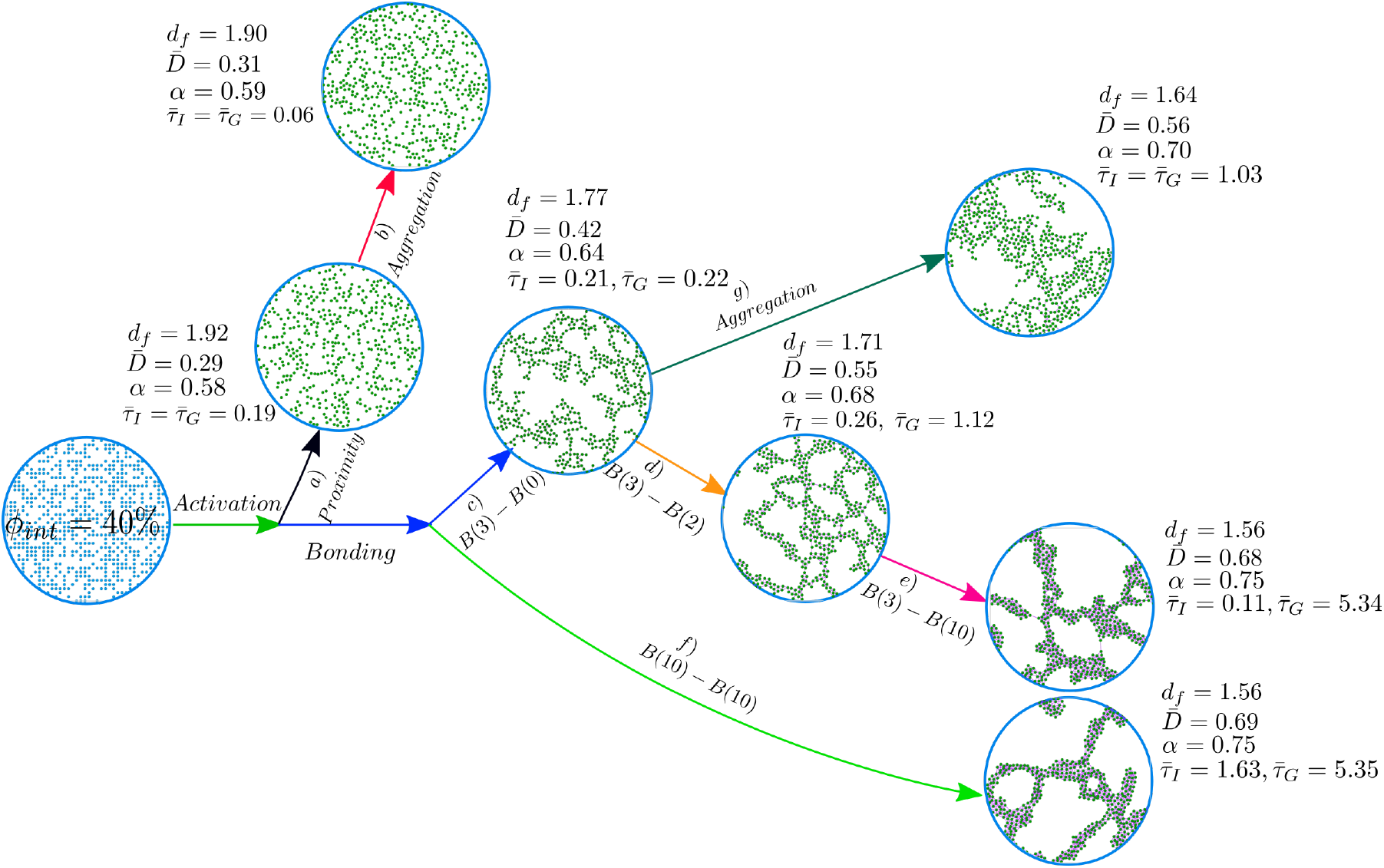
A biomarker diagram is provided step by step regarding the implementation of different mechanisms in our model for *ϕ_int_* = 40%. Biomarkers include *d_f_, D*_∞_, *α, τ_G_* and *τ_G_*.

The results show how altered particle interactions and initial concentrations lead to distinct changes in cluster morphology and kinetics. The method is capable of constructing a wide range of microstructures, with a *d_f_* range of 1.4±0.01 to 2±0.01. Furthermore, it is obvious that the use of a sole biomarker, such as *d_f_*, is insufficient to provide a thorough differentiation between clusters. We proposed that additional characterization of the mean square displacement of solvent particles (or probes) confined within the clusters provides an integral characterization of biological clusters. Furthermore, by analyzing the time-dependent diffusion coefficient or anomalous diffusion exponent, we were able to distinguish two physically distinct regimes: the short-time regime and the long-time regime approaching asymptotic Gaussian diffusion. This approach highlighted essential biological system features that were not evident from *d_f_* alone.

The introduced GMC has the potential to further study complex cluster mechanisms, such as those related to blood coagulation, and could range from fibrin network formation to platelet aggregation. Future studies will examine the mechanical properties of different cluster configurations, which frequently have biological implications. For example, changes in the mechanical properties of blood clots have been observed in coronary artery disease and venous thrombosis, highlighting the potential of using viscoelastic measurements to improve our understanding of these conditions.

## Supplementary Information

An online supplement to this article can be found by visiting BJ Online at http://www.biophysj.org.

## Author Contributions

N.M and M.E conceived and supervised the project. E.Z designed the computational experiments and conducted the simulations and data processing. The original draft of the manuscript was written by E.Z and N.M. and all authors contributed to the final version of the manuscript. N.M. developed the numerical implementation. All the authors discussed and analyzed the results. All authors approved the final version of the manuscript.

## Acknowledgments

This research is supported by Basque Business Development Agency under ELKARTEK 2019 programme (bmG19 project: grant KK-2019/00015) and through the “Mathematical Modeling Applied to Health” Project. Also by the Basque Government through the BERC 2022-2025 program and by the Ministry of Science and Innovation: BCAM Severo Ochoa accreditation CEX2021-001142-S / MICIN / AEI / 10.13039/501100011033. N.M acknowledges the support from the European Union’s Horizon 2020 under the Marie Sklodowska-Curie Individual Fellowships grant 101021893, with acronym ViBRheo.

