## Supplementary Information for "Computational Mesoscale Framework for Biological Clustering and Fractal Aggregation"

### 1 Supplementary Equations

The thermal fluctuation is included in the model by

$$m\frac{\mathrm{d}\tilde{\mathbf{v}}_i}{\mathrm{d}t} = \sum_j \overbrace{\left(A_{ij}\mathrm{d}\overline{\mathbf{W}}_{ij} + B_{ij}\frac{1}{D}\mathrm{tr}[\mathrm{d}\mathbf{W}_{ij}]\right)\cdot \mathbf{e}_{ij}}^{\mathbf{f}^{\mathrm{r}}_{ij}}, \tag{S1}$$

where $\overline{\mathbf{W}}_{ij}$ is a matrix of independent increments of a Wiener process for each pair $i, j$ of particles, and $\overline{\mathbf{W}}_{ij}$ is its traceless symmetric part, given by

$$\mathrm{d}\overline{\mathbf{W}}_{ij} = \frac{1}{2}\left[d\mathbf{W}_{ij} + \mathrm{d}\mathbf{W}^T_{ij}\right] - \frac{\delta^{\alpha\beta}}{D}\mathrm{tr}[\mathrm{d}\mathbf{W}_{ij}],$$

where $D$ is the dimensionality of the system. To satisfy the fluctuation-dissipation balance the amplitude of the thermal noises $A_{ij}$ and $B_{ij}$ are related to the friction coefficients $a_{ij}$ and $b_{ij}$ through

$$A_{ij} = \left[4k_BTa_{ij}\frac{F_{ij}}{\rho_i\rho_j}\right]^{1/2}, \tag{S2}$$

$$B_{ij} = \left[4k_BT\left(b_{ij} - a_{ij}\frac{D-2}{D}\right)\frac{F_{ij}}{\rho_i\rho_j}\right]^{1/2}, \tag{S3}$$

To describe the variation of the pressure with the density of the system we adopt the Cole equation (a.k.a. Tait's equation of state) given by $p_i = c^2\rho_0/7\left[(\rho_i/\rho_0)^7 - 1\right] + p_b$ (S4) where $c$ is the speed of sound on the fluid, and $\rho_0$ is the reference density. The term $c^2\rho_0/7$ corresponds to the reference pressure of the system, given by $c^2 = \partial p/\partial\rho|_{\rho=\rho_0}$. The parameter $p_b$ is a background pressure, that provides numerical stability by keeping the pressure of the system always positive.

For the interpolant function, we adopt the Lucy kernel [31] typically used in SDPD

$$W(r) = \begin{cases} \frac{w_0}{h^D}\left(1+\frac{3r}{h}\right)\left(1-\frac{r}{h}\right)^3, & r/h < 1. \\ 0, & r/h > 1, \end{cases} \tag{S5}$$

where $w_0 = 5/\pi$ or $w_0 = 105/16\pi$ for two or three dimensions, respectively. For a comprehensive description of the SDPD method, the reader is referred to [75].

### 2 MSD calculation

The time-averaged MSD is calculated from the trajectory of moving $(\mathbf{S})$ particles in the range $t = 0, ..., T$.

$$\langle\delta^2(\Delta)\rangle = \frac{1}{T-\Delta}\int_0^{T-\Delta}[\mathbf{r}(t+\Delta) - \mathbf{r}(t)]^2 dt \tag{S6}$$

where $\mathit{\Delta}$ is the so-called lag time, which defines the size of a window slid along the trajectory $\mathbf{r}(\mathrm{t})$. The trajectory length $T$ is also referred to as measurement time. Besides the individual time traces $\langle\delta^2(\mathit{\Delta})\rangle$ we also considered the average over $N$ individual trajectories:

$$\langle\langle\delta^2(\Delta)\rangle\rangle = \frac{1}{N}\sum_{k=1}^{N}\langle\delta^2(\Delta)\rangle \tag{S7}$$

In normal diffusion, $\langle\langle\delta^2(\Delta)\rangle\rangle \sim Dt$, where the diffusion coefficient $D$ is constant. In pure anomalous subdiffusion, $\langle\langle\delta^2(\Delta)\rangle\rangle \sim t^\alpha$, $\alpha < 1$ at all times, where $\alpha$ is the anomalous diffusion exponent. The diffusion coefficient is, therefore, time-dependent, $D(t) \sim 1/t^{1-\alpha}$, appropriately modified to give the proper limit at $t = 0$, say

$D(t) = D_0/(1 + t^{1-\alpha})$. The case of interest here is transient anomalous subdiffusion, in which there is a crossover from anomalous subdiffusion at short times to normal diffusion at long times,

$$\langle\langle \delta^2(\Delta) \rangle\rangle \sim \begin{cases} t^\alpha & \text{for } t \ll t_{CR2} \\ t & \text{for } t \gg t_{CR2} \end{cases} \tag{S8}$$

where $t_{CR2}$ is the crossover time.

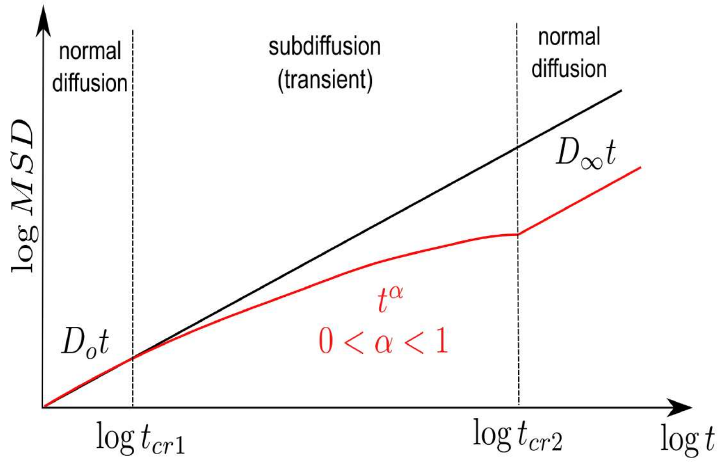


Figure S1: Behavior of MSD in the complex clusters

#### 3 Supplementary Results

paths.png paths.png

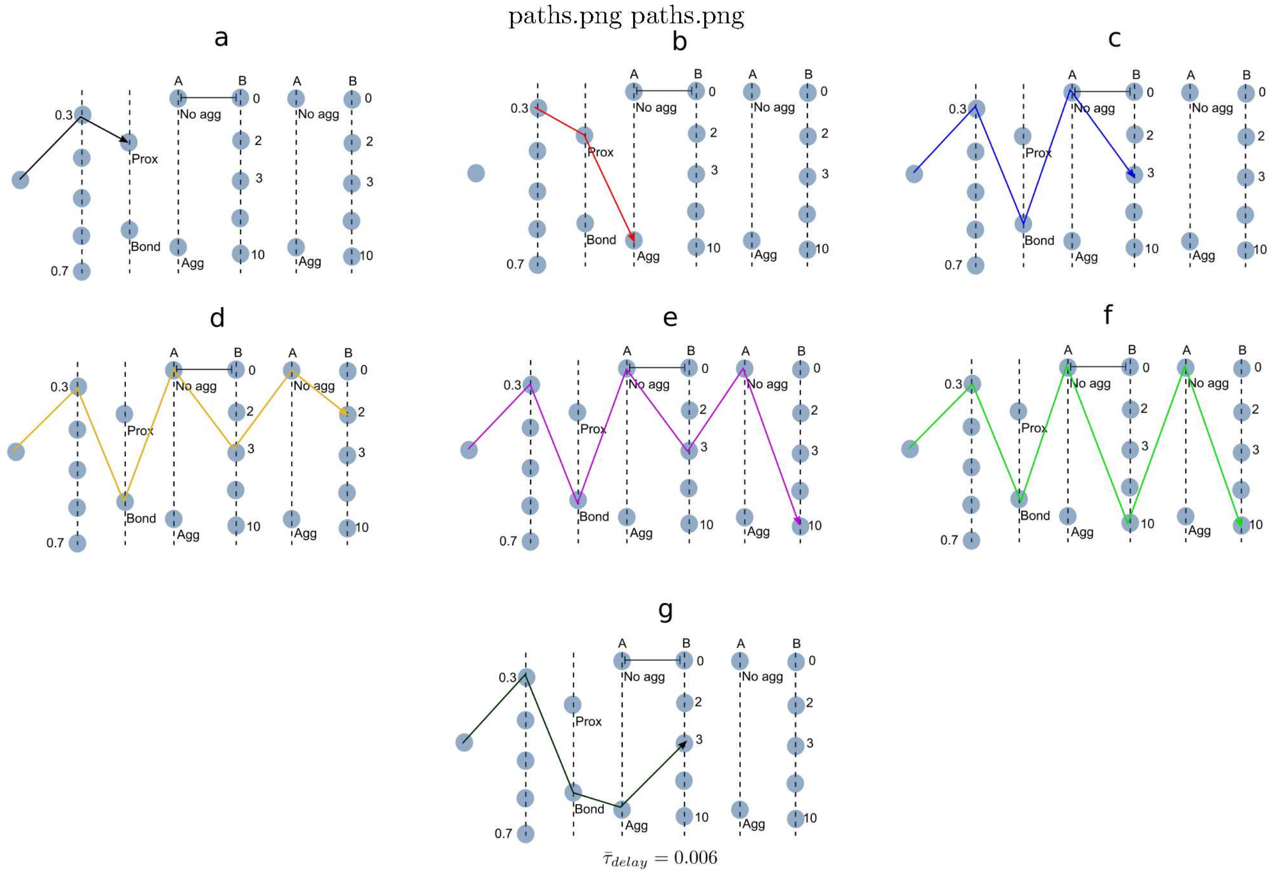


Figure S2: The complete set of mechanisms (**a**)–(**g**) paths

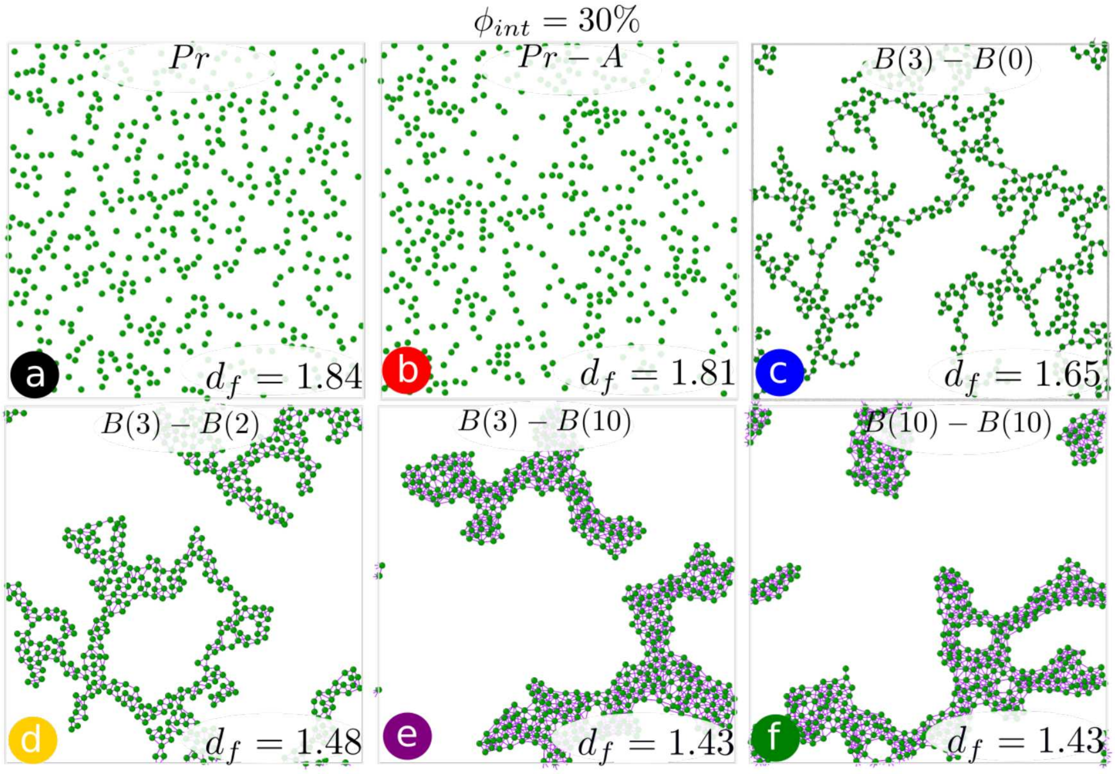


Figure S3: Final cluster morphology and $d_f$ are shown for six alternative mechanisms $(a, b, c, d, e, f)$ and $\phi_{int} = 30\%$

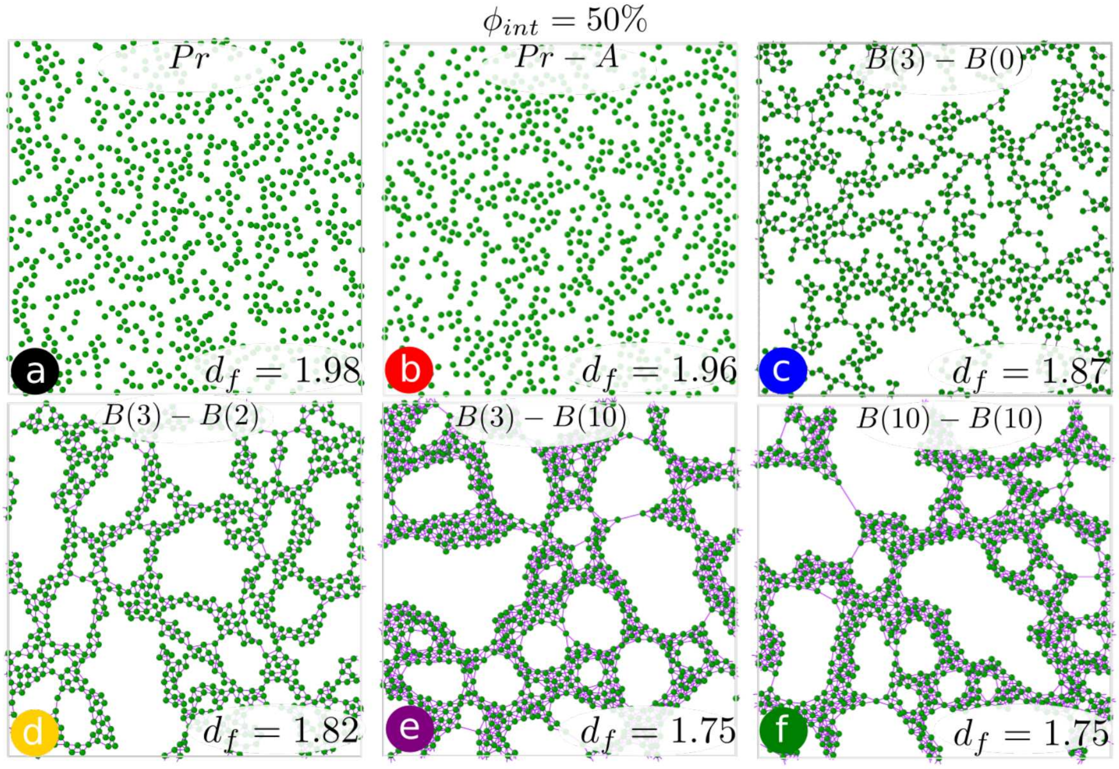


Figure S4: Final cluster morphology and $d_f$ are shown for six alternative mechanisms $(a, b, c, d, e, f)$ and $\phi_{int} = 50\%$

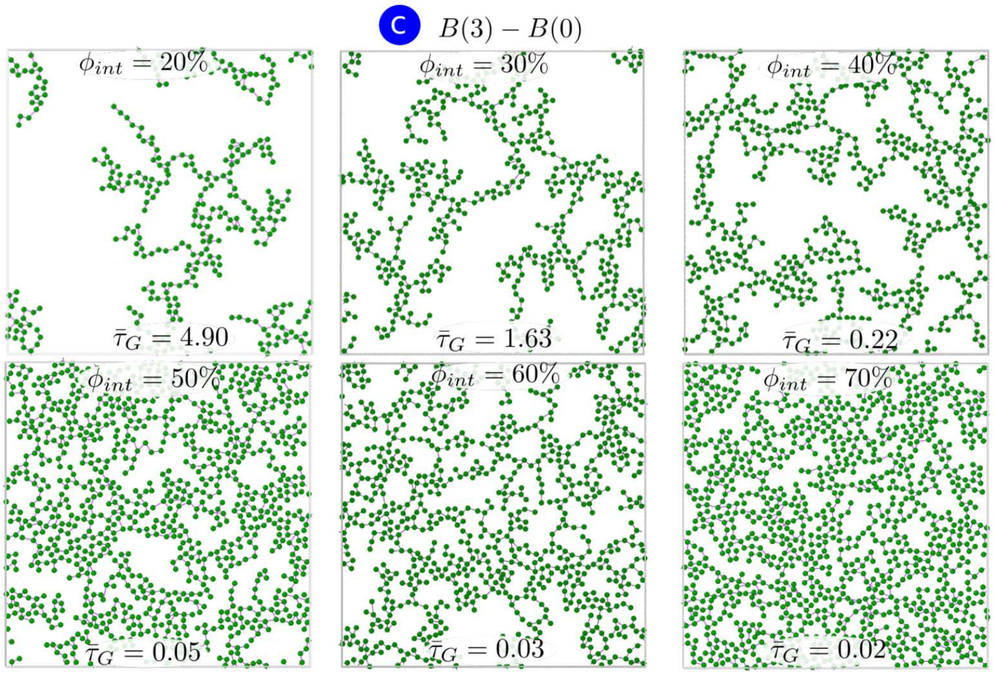


Figure S5: Final cluster morphology and $\bar{\tau}_G$ are shown for six alternative concentrations $\phi_{int}$ = 20%, 30%, 40%, 50%, 60% and mechanism $c$ ($B(3)\_B(0)$)

#### 3.1 Effect of first-stage bonding number

We explore the morphology of clusters for three distinct mechanisms where the second-stage bonding or consolidation stage is off ($m = 0$), in order to examine the effect of the first-stage bond number. We do this by altering $n$ bonds number created at the initial stage of bonding for $n = 2, 3, 10$ (mechanisms: $B(2)\_B(0), B(3)\_B(0), B(10)\_B(0)$) and three different concentrations of $\phi_{int} = 30\%, 40\%, 50\%$ as shown in Fig. S6. We define the parameter $N_B/N_A$ (the average number of bonds formed between (**A**) particles) as a biomarker for studying the cluster structure. One might observe that $N_B/N_A$ does not exceed 1 for all mechanisms and there is no difference between the structure of clusters for $n > 2$. However, because there are only two allowed bonds in the mechanism $B(2)\_B(0)$, the cluster structure resembles a chain, and the activation time for every (**P**) particle is quite long, especially for lower concentration values. There are still (**P**) particles in the system at all three concentrations of this mechanism as depicted in Fig. S6. It is possible to deduce that the characterization of a cluster is unaffected by the number of bonds in the first-stage bonding for $n > 2$. However, regardless of how large $n$ is, $N_B/N_A$ is equal 1. This means that initial stage bonding can only resemble immature clots.

#### 3.2 Effect of second-stage bonding number

To investigate the effect of the second-stage bond number, the first-stage bonding number, $n$ is fixed to 3, and the bond number in the second stage, $m$ is adjusted to $m = 0, 2$, and 10, equal to mechanism types $c$, $d$ and $e$ ($B(3)\_B(0)$, $B(3)\_B(2)$ and $B(3)\_B(10)$). In Fig. S7 the cluster morphologies of these three mechanisms and different concentrations $\phi_{int} = 30\%, 40\%, 50\%$ are shown. When there is no bond in the second stage ($m = 0$), the average number of bonds formed between (**A**) particles, $N_B/N_A = 1$ whereas for $m = 2$ and $m = 10$, the cluster structure becomes more branched, solid and stable with $N_B/N_A > 1$, resulting the lower value of $d_f$. It implies that we can only imitate mature clots using second-stage bonding. However, it appears that the second-stage bond number has altered not only the fractality of the cluster but also the cluster kinetic characteristics, as evaluated by MSD measurement (outlined in this study as a new biomarker).

#### 3.3 Effect of delay time

This section's primary goal is to examine the impact of the delay time between the aggregation process and the first-stage bonding on the cluster morphology and $\tau_I$. We consider mechanism $g$ ($B(3)A_{ps}\_B(0)$) and $\phi_{int} = 40\%$ to see this effect. The delay time is $\bar{\tau}_{delay} = 0.006$ that is dimensionless with $\tau_{diff}$. In this manner, we apply initiation bonding for every $\tau_{delay}$ duration of aggregation. The results in Fig. S8.$c$ show that by delaying the time between bonding and aggregation, $\bar{\tau}_I$ increases in comparison to the case $\tau_{delay} = 0$ (seen in

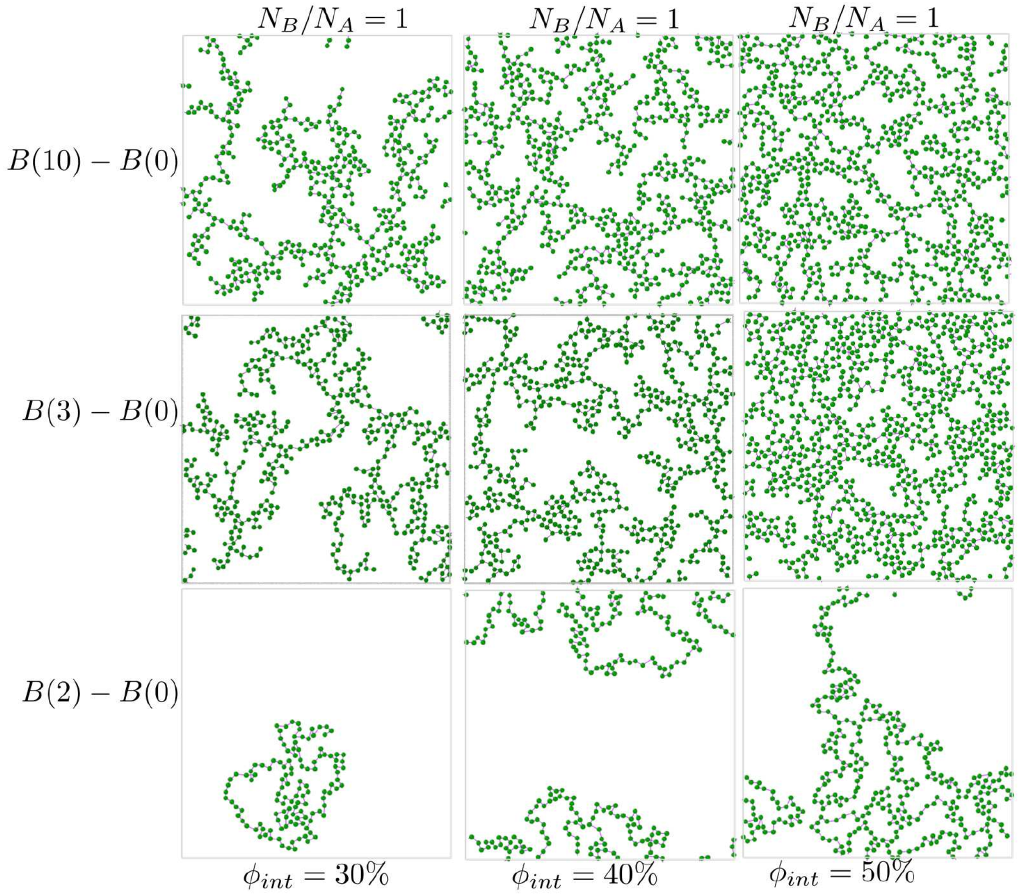


Figure S6: Effect of the first-stage bond numbers on cluster morphology for various mechanisms: $B(2)\_B(0), B(3)\_B(0), B(10)\_B(0)$ and $\phi_{int} = 30\%, 40\%, 50\%$. The average number of bonds formed between (**A**) particles ($N_B/N_A$) are shown.

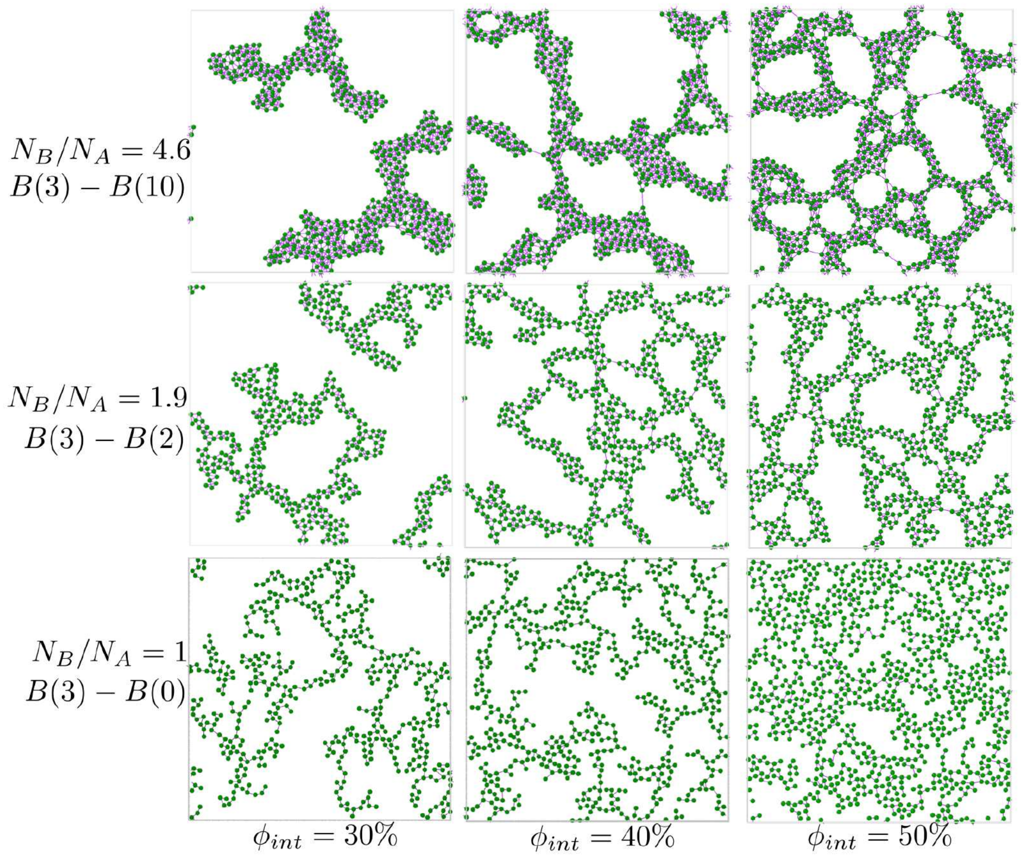


Figure S7: Effect of the second-stage bond numbers on cluster morphology for various mechanisms of $(c, d, e)$ and $\phi_{int} = 30\%, 40\%, 50\%$. The average number of bonds formed between (**A**) particles ($N_B/N_A$) are shown.

Fig. S8.*b*) also $d_f$ and final cluster morphology profoundly changes. We also infer that adding aggregation to mechanism $c$ ($B(3)\_B(0)$) without delay time (seen in Fig. S8. *a*, *b*) has little effect on $d_f$ and $\tau_I$. In Fig. S9, which depicts cluster evolution with time for $\bar{\tau} = 0.006$, can observe how the droplet form of (**P**) particles (because of aggregation) makes the first-bonding stage slower. Delay time, replicates the activation delay time for platelets during the clot-forming process. It is important because a delay in the activation time for platelets can increase the risk of bleeding, while an overactive response can lead to an increased risk of blood clots and related health problems.

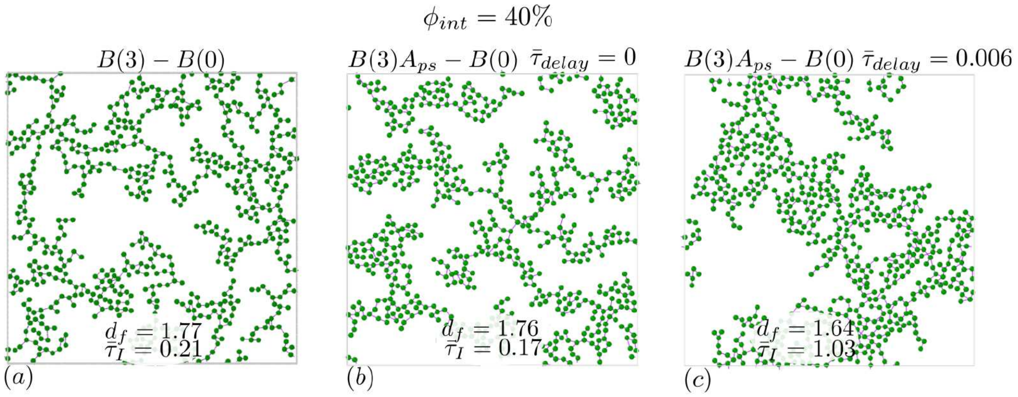


Figure S8: Effect of delay time between first-stage bonding and aggregation on cluster structure. The final morphology, $d_f$ and $\bar{\tau}_I$ are displayed for $\phi_{int} = 40\%$ and for a)mechanism $c$, b) mechanism $g$, and $\bar{\tau}_{delay} = 0$, c) mechanism $g$, and $\bar{\tau}_{delay} = 0.006$ to see how the delay time between aggregation and activation change cluster structure.

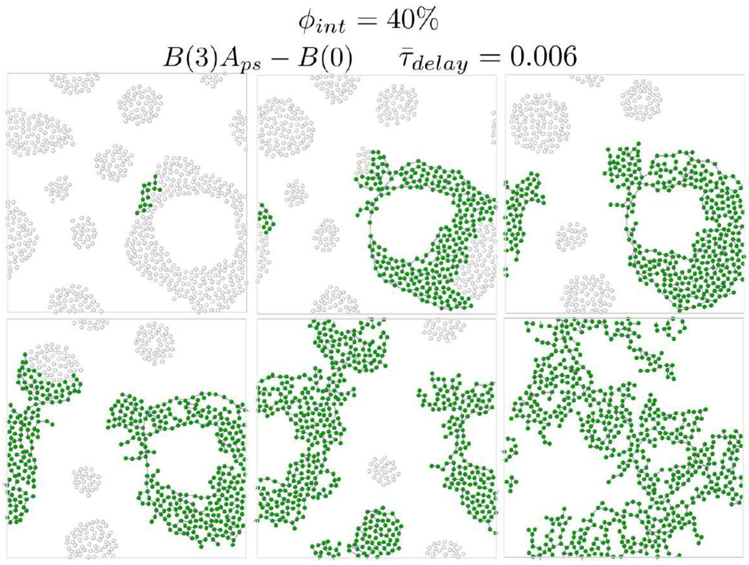


Figure S9: Cluster evolution with time for mechanism $g$ and $\bar{\tau}_{delay} = 0.006$ of first-stage bonding and aggregation. The activation of passive particles can also be seen in the step-by-step morphology of this mechanism.

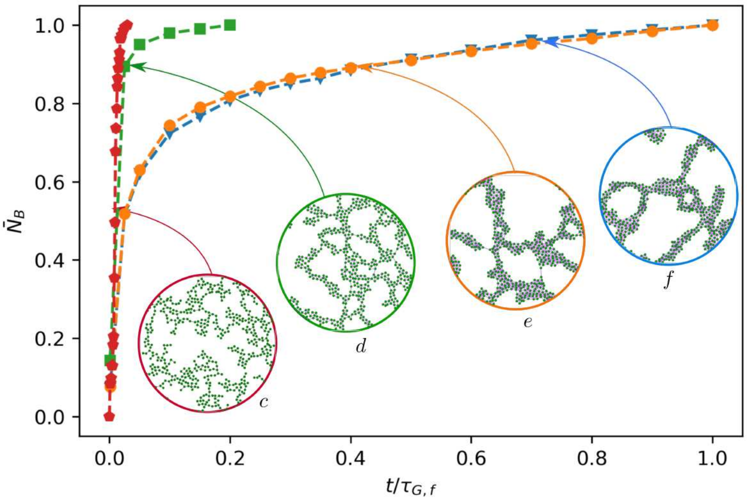


Figure S10: Bonds number evolution with time for various mechanisms of $c, d, e, f$ and $\phi_{int} = 40\%$. Each curve is dimensionless with its values in a steady state or gelling state, and time is also dimensionless with $\tau_G$ of mechanism $f$. The final morphology for each mechanism is displayed.

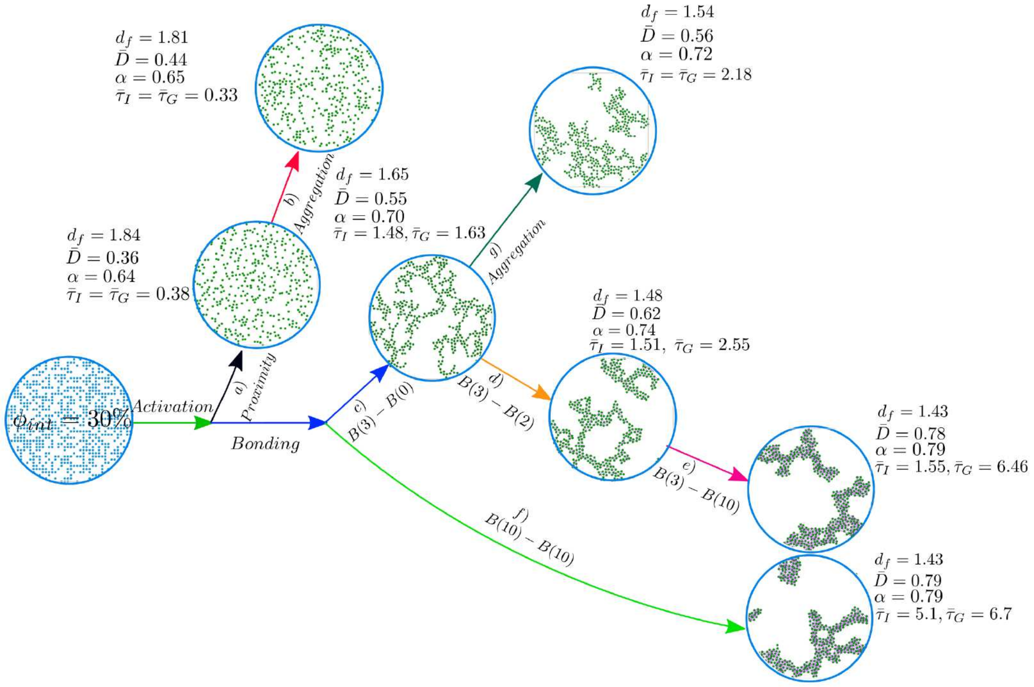


Figure S11: A biomarker diagram is provided step by step regarding the implementation of different mechanisms in our model for $\phi_{int} = 30\%$. Biomarkers include $d_f$, $D_\infty$, $\alpha$, $\tau_I$ and $\tau_G$.

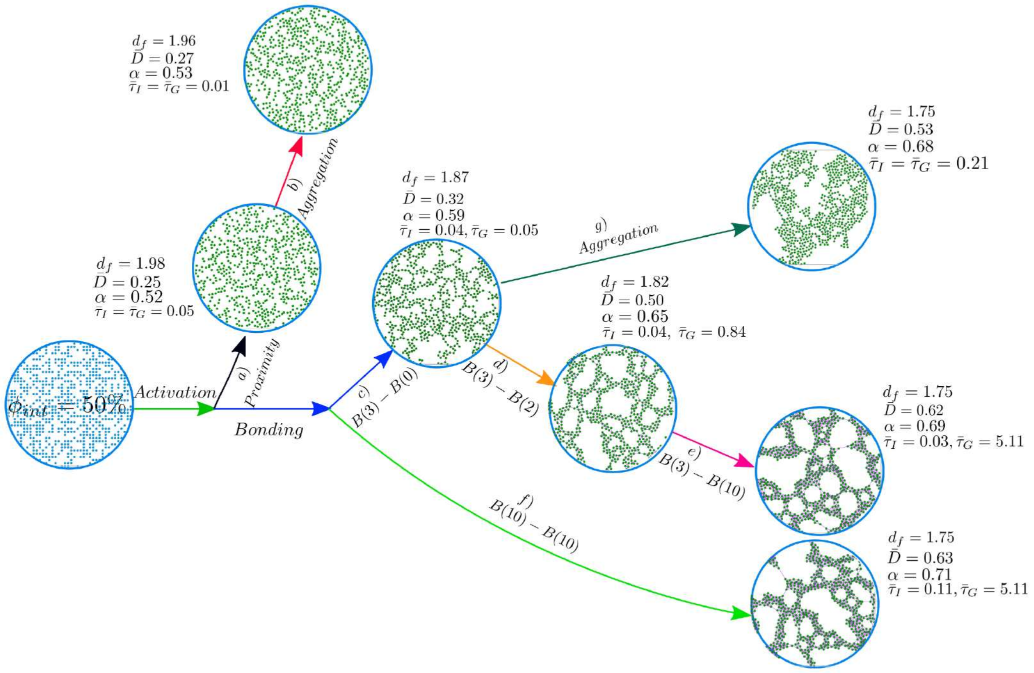


Figure S12: A biomarker diagram is provided step by step regarding the implementation of different mechanisms in our model for $\phi_{int} = 50\%$. Biomarkers include $d_f$, $D_\infty$, $\alpha$, $\tau_I$ and $\tau_G$.

### 4 Supplementary Tables

Table S1: Input parameters of the SDPD method

| | |
|---|---|
| Domain size $[L_x \times L_y]$ | $(40dx)\times(40dx)$ |
| Total number of particles $(N_t)$ | 1600 |
| Mass$(m)$ | 0.04 |
| viscosity$(\eta)$ | 10 |
| $k_B T$ | 0.1 |
| Density$(\rho_0)$ | 1 |
| Pressure$(p_0)$ | 50 |
| Speed of sound $(c_s)$ | 40 |
| Time step $(dt)$ | $10^{-4}$ |
| Initial lattice spacing$(dx)$ | 0.2 |
| Cutoff radius $(h)$ | $4dx$ |

Table S2: Input parameters of the bond and surface tension potentials

| Bond Strength, $D$ (Eq. 7) | Stiffness Parameter, $\alpha$ (Eq. 7) | Bond Length, $r_0$ (Eq. 7) | Surface Tension, $\sigma$ (Eq. 6) |
|---|---|---|---|
| 30 | 1 | 0.2 | 0.5 |

Table S3: $d_f$ values (with the standard deviation equal to $\pm 0.01$) for different, $\phi_{int}$ and six different mechanisms

| Mechanisms | | | Concentrations |
|---|---|---|---|
| | $\phi_{int} = 30\%$ | $\phi_{int} = 40\%$ | $\phi_{int} = 50\%$ |
| $a$ | $d_f = 1.84$ | $d_f = 1.92$ | $d_f = 1.98$ |
| $b$ | $d_f = 1.81$ | $d_f = 1.90$ | $d_f = 1.96$ |
| $c$ | $d_f = 1.65$ | $d_f = 1.77$ | $d_f = 1.87$ |
| $d$ | $d_f = 1.48$ | $d_f = 1.71$ | $d_f = 1.82$ |
| $e$ | $d_f = 1.43$ | $d_f = 1.56$ | $d_f = 1.75$ |
| $f$ | $d_f = 1.43$ | $d_f = 1.56$ | $d_f = 1.75$ |
